# Molecular heterogeneity of quiescent melanocyte stem cells revealed by single-cell RNA-sequencing

**DOI:** 10.1101/2023.12.19.571712

**Authors:** Joseph W. Palmer, Nilesh Kumar, Luye An, Andrew C. White, M. Shahid Mukhtar, Melissa L. Harris

## Abstract

Melanocyte stem cells (McSCs) of the hair follicle are a rare cell population within the skin and are notably underrepresented in whole-skin, single-cell RNA sequencing (scRNA-seq) datasets. Using a cell enrichment strategy to isolate KIT+/CD45-cells from the telogen skin of adult female C57BL/6J mice, we evaluated the transcriptional landscape of quiescent McSCs (qMcSCs) at high resolution. Through this evaluation, we confirmed existing molecular signatures for qMcCS subpopulations (e.g., *Kit+, Cd34+/-*, *Plp1+, Cd274+/-, Thy1+, Cdh3+/-*) and identified novel qMcSC subpopulations, including two that differentially regulate their immune privilege status. Within qMcSC subpopulations, we also predicted melanocyte differentiation potential, neural crest potential, and quiescence depth. Taken together, the results demonstrate that the qMcSC population is heterogenous and future studies focused on investigating changes in qMcSCs should consider changes in subpopulation composition.

**Significance:** Single cell transcriptomics has revolutionized our ability to interrogate the dynamic nature of tissues. Here we provide a high-resolution map of the melanocyte stem cell population during quiescence. This map provides one of few examples highlighting broad heterogeneity in stem cells during the quiescent cell state. The map also unifies previous observations using other cell, molecular and functional analyses to define the unique features of the quiescent melanocyte stem cell population. This data provides a valuable resource to individuals interested in further evaluating aspects of cellular quiescence in stem cells broadly or melanocyte stem cells specifically.

## Introduction

Melanocyte stem cells (McSCs) are a rare stem cell population that accounts for a very small percentage of the skin, representing ∼1-2% of the total cells of the dermis ^1,2^. Prior to the advent of single-cell transcriptomic methods, researchers relied on bulk tissue methods to investigate gene expression differences in McSCs under various genetic or external perturbations. These studies provide valuable insights into the molecular regulation of McSC function during hair repigmentation and in melanocyte disease ^1–4^. Bulk sequencing, however, lacks the ability to resolve gene expression differences between subpopulations of McSCs and may mask outcomes where certain McSC groups are lost or gained. These events are instead inappropriately interpreted as population-wide global transcriptional change.

The concept of McSC heterogeneity is supported by observations that the McSC population is comprised of subpopulations that are functionally, molecularly, and anatomically distinguishable. For example, using mini-chromosome maintenance protein (Mcm) as a marker of replication in mice, only about half of McSCs become proliferative (Mcm+) during the hair growth stage of the hair cycle, and these McSCs rely on KIT signaling for their survival and are radioresistant. The remaining 50% are Mcm– and appear quiescent, KIT-independent, yet radiosensitive ^5^. McSCs reside within the stem cell niche of the hair follicle ^6^. During hair dormancy McSCs localize to two distinct regions, the bulge and secondary hair germ (sub-bulge), and these regions can be defined molecularly by CD34 and P-cadherin (CDH3) expression, respectively ^7^. These markers also specify the McSCs that exist within these two spaces, which further translates to differences in their cell potential. CD34+ McSCs found in the hair bulge represent a less differentiated multipotent population that retains the ability to differentiate into myelinating Schwann cells or pigmenting melanocytes under particular conditions ^2^. More recently, we described a novel subpopulation of quiescent McSCs (qMcSCs) that express the immune checkpoint protein, PD-L1, that is retained with age ^8^. Based on these observations, it appears that not all McSCs are made to be equal. Understanding the differences between McSC subpopulations and how they interrelate will shed light on how each subpopulation might contribute to the maintenance of physiological pigmentation and potentially highlight those subpopulations prone to disease.

In mouse, early attempts to characterize individual McSCs preceded next-generation sequencing technologies and, while limited in the number of cells and transcripts evaluated, hinted at the possibility of variability within the McSC population based on the differential expression of melanogenic genes ^9^. More recent single-cell RNA sequencing datasets of McSCs confirm this possibility revealing clear differences in the McSC pool when isolated from hairs during dormancy (qMcSCs) or activation (activated McSCs; aMcSCs) and revealed two identifiable clusters of aMcSCs ^10,11^. However, these studies fail to capture the complexity of the qMcSCs subpopulations predicted by the functional, molecular, and anatomical differences mentioned above, likely due to a limited number of qMcSCs evaluated (Infarinato et al. 2020 = 104 qMcSCs; Joost et al. 2020 = 5 qMcSCs). In zebrafish, single-cell profiling reveals two distinct populations of *mitfa-*expressing qMcSCs defined by *aox5* low or high expression. These two populations also respond uniquely during pigment regeneration with *mitfa^+^aox5^lo^* qMcSCs predicted to give rise to melanocytes via direct differentiation and *mitfa^+^aox5^hi^*qMcSCs giving rise to melanocytes through a cycling precursor ^12^. Altogether, these observations beg the question, just how different are qMcSCs?

Here we present in high-resolution the transcriptional landscape of mouse qMcSCs. Our analysis helps to coalesce previously identified qMcSC subpopulations, reveals novel qMcSC subpopulations, highlights the key gene expression differences between these subpopulations, and shows that qMcSCs can be mapped along a differentiation trajectory. Using predictive modeling we show a range of quiescence ‘depth’ across our cell clusters with the more differentiated qMcSCs associated with a shallower quiescent state. Finally, we take a focused look at the two qMcSCs supopulations defined by *Cd274* expression and make the novel prediction that *Cd274* may be an alternate mechanism to provide immune privilege to stem cells that retain high potential for antigen presentation (MHC Class I). Altogether this study presents the first comprehensive overview of the qMcSC population of the hair follicle during dormancy and extends our understanding of the heterogeneity observed in this rare stem cell population.

## Results

### Analysis of qMcSCs from scRNA-seq of Whole Dermis

Our approach for evaluating qMcSCs was twofold. First, we considered qMcSCs in the context of the dermal tissue in which they exist, and second, we used an enrichment strategy for qMcSCs in order to increase the resolution of our analysis (Figure 1a). In both contexts, the dermis of ∼10-week-old, female, C57BL/6J mice is separated from the epidermis and dissociated into single cells. This single-cell suspension was used directly for ‘whole dermis’ scRNA-seq (n=1), or further enriched for KIT+/CD45-qMcSCs prior to scRNA-seq using our previously validated flow cytometry method (n=3) ^1,8^.

**Figure 1:**
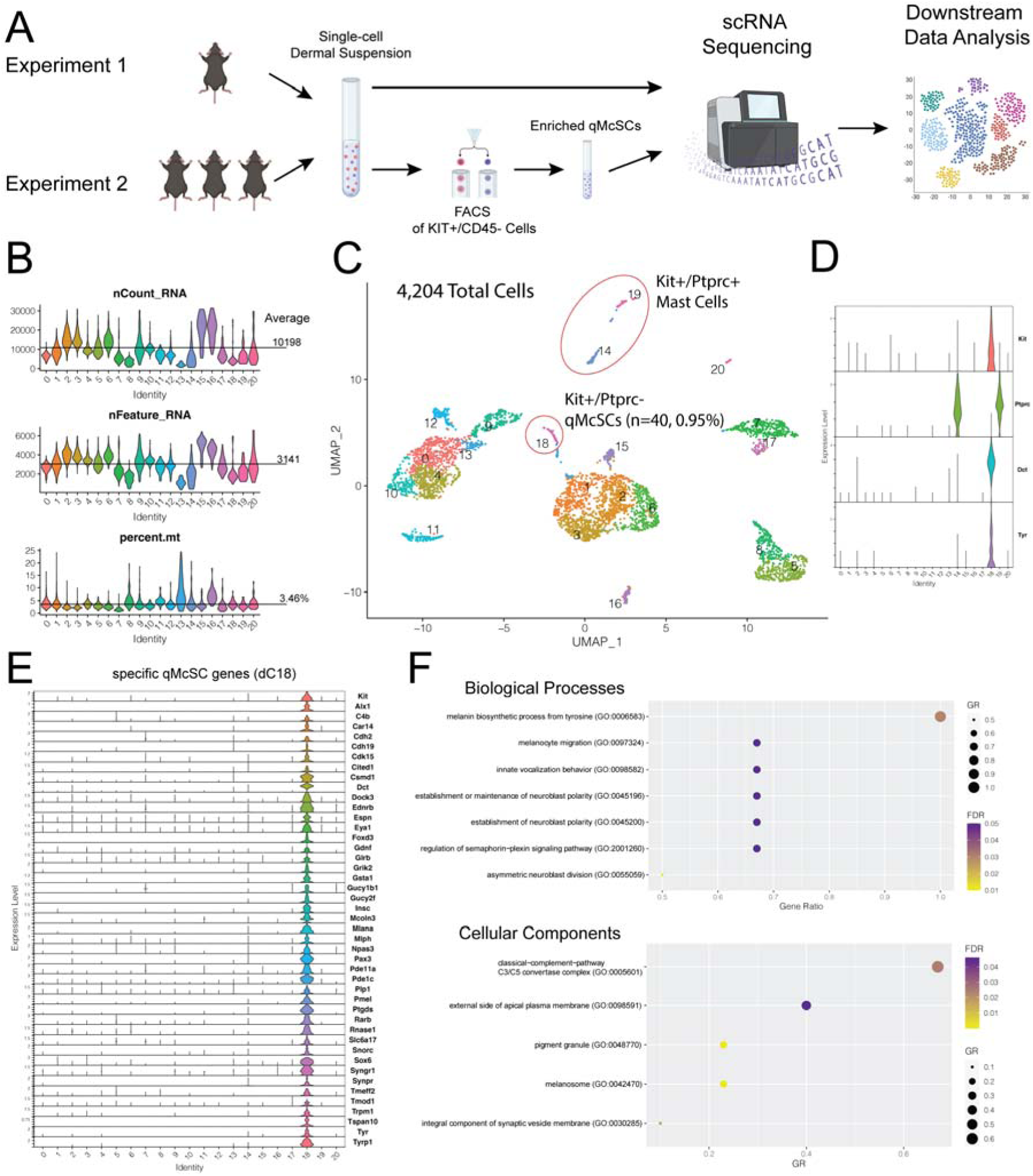
Diagram of experimental design and identification of markers specific to qMcSCs in the dermis. (A) Diagram of experimental design used in this study. (B) Overview of the quality control metrics of dermal cells showing the average RNA, feature counts and percent mitochondrial genes detected across clusters. (C) UMAP of dermal cells arranged into 21 dermal clusters (dCs). Circled are the *Kit+/Cd45(Ptprc)-* qMcSCs (dC18), a rare population (0.95%) of cells with the telogen dermis. Also circled are the *Kit+/Cd45(Ptprc)+* mast cells (dC14, dC19) that are negatively selected away during the cell enrichment step of Experiment 2, shown in 1A. (D) Violin plots of key genes used to identify qMcSCs. (E) Violin plot showing the 45 global markers that are unique to the dC18 qMcSC cluster. (F) Enrichment analysis of select biological processes and cellular components using the 177 upregulated DEGs expressed by the qMcSC dC18.

To get a sense of how qMcSCs compare with their neighboring dermal cells we first analyzed the whole dermis. Following standard quality control methods to filter and normalize the data, evaluation of the whole dermis resulted in a total of 4,204 cells resolved into 21 clusters (resolution = 1.0) with an average of 10198 nCounts, 3141 nFeatures, and 3.5% mitochondria genes detected across cells (Figure1b-c). Based on the localized expression of melanocyte markers *Kit*+/*Dct*+/*Tyr*+ and little to no cells expressing *Ptprc,* the gene encoding the mast cell marker CD45, we assigned dermal cluster 18 (dC18, n = 40 cells) as the qMcSC population (Figure 1c-d). dC18 roughly amounts to 1% of dermal cells, an amount that matches previous measurements of qMcSC abundance in the dermis using an alternate gene reporter method ^13^. *Kit* is also expressed by mast cells, yet the clusters with *Ptprc+* cells, dC9, and dC14, have an average *Kit* expression of 0.11 and 0.08, respectively. This level of *Kit* expression in these *Ptprc+* cells is substantially higher than the 0.012 average background *Kit* expression calculated across all other cell clusters yet notably lower than the average *Kit* expression of 1.31 detected in qMcSCs of dC18. These data support the conclusion that *Kit+* qMcSCs in dC18 can be uniquely distinguished from other *Ptprc+/Kit+* mast cell clusters.

Global differential analysis (min.pct = 0.1, logfc.threshold = 0.25) comparing dC18 to all other clusters resulted in 177 differentially upregulated genes (DEGs; q-value > 0.05, avg.L2FC > 0.25; Supplemental File 1). Of these 177 upregulated DEGs, 45 were specific to dC18, meaning not observed at a high level of expression in a significant number of cells in any of the other clusters (Figure 1e). Included in these specific genes were 8 transcriptions factors (TFs): *Foxd3*, *Pax3*, and *Sox10* have established roles in the development of melanoblasts from the neural crest, *Sox6* has been linked to melanogenesis in Alpaca ^14^, and *Alx1*, *Npas3*, *Plagl1*, *Rarb* with yet unknown functions in pigmentation. Cells within the dC18 cluster also express specific genes involved in melanocyte signaling and melanin biosynthesis, *Kit*, *Dct*, *Ednrb*, *Tyr*, and *Tyrp1* ^15^, along with 13 genes known to cause pigmentation defects in humans or mice or are otherwise involved in pigmentation including *Cdh2*, *Cited1*, *Mcoln3*, *Mlph*, *Pdella*, and *Slc24a3*^16^. Lastly, we find additional genes to be specific to the qMcSC cluster including some that may confer benefits to a quiescent stem cell population. This includes the Schwann cell precursor marker *Cdh19*, which has also been observed as a unique marker for melanocytes in human skin ^17,18^. We also observed *Cdk15*, a protein that can negatively regulate TRAIL-induced apoptosis, along with the double and single-strand degrading ribonuclease *Rnase1*, which can promote stemness in breast cancer cells ^19–24^.

Gene set enrichment analysis (GSEA) of the 177 upregulated DEGS from dC18 showed biological processes such as melanin biosynthetic process from tyrosine (FDR = 3.24E-02, Gene ratio = 1.0), regulation of semaphorin-plexin pathway (FDR = 4.82E-02, Gene ratio = 0.67), the establishment of neuroblast polarity (FDR = 4.79E-02, Gene ratio = 0.67), and asymmetric neuroblast division (FDR = 5.72E-03, Gene ratio = 0.5) associated with genes with increased expression in the dC18 cluster (Figure 1f, Supplemental File 1). These findings further confirm that dC18 can be identified as qMcSCs with qualities of both melanocytes (pigmentation) and stem cells (polarity and asymmetric division). Additionally, we find that certain cellular components are also enriched in this cell cluster including the classical-complement-pathway C3/C5 convertase complex (*C2* and *C4b*), external side of the apical plasma membrane (*Slc7a5*, *Slc38a1*), melanosome and pigment granule (*Sytl2*, *Mlph*, *Tyrp1*, *Mlana*, *Pmel*, *Tyr*, *Dct*, and *Myo5a*), and genes associated with an integral component of synaptic vesicle membrane (*Syngr1*, *Slc6a17*, *Atp6v0a1*, and *Synpr*) (Figure 1g, Supplemental File 1). Lastly, dC18 also upregulates the DEGs *Cd274* and *Plp1*, two genes previously associated with McSCs and the melanocyte lineage. Our group showed that a subpopulation of qMcSCs expresses PD-L1, the protein product of *Cd274*, during the telogen stage of the hair cycle ^8^. Others showed that PLP1 is expressed by Schwann cell precursors that contribute to extracutaneous melanocytes of the heart, inner ear, meninges, and skin ^25^.

Taken together we show that the telogen dermis contains a distinct population of qMcSCs that are identifiable by the upregulation of several unique genes. With future validation, these markers will serve as useful tools to aid in McSC identification and experimental comparison. These qMcSCs also express markers of subpopulations that suggest additional compartmentalization within the qMcSC pool and warrant further evaluation using enriched qMcSC scRNA-seq samples to better resolve these subclusters. However, given the relatively low abundance of cells that comprise dC18 (n=40) the resolution required to further delineate these known qMcSC subpopulations or other novel subpopulations requires additional McSC enrichment.

### Generation of High-Resolution scRNA Transcriptomic Map of KIT+/CD45-qMcSCs

To overcome the limitations associated with investigating qMcSCs in datasets generated from the whole dermis, we enriched KIT+/CD45-qMcSCs as mentioned above (Figure 1a). Standard quality control methods were used to filter poor-quality cells. Clusters with *Kit* expression comparable to the *Kit*-negative clusters observed in the dermis (average cluster expression < 0.1) were also removed. The remaining cells were clustered using UMAP into 8 major clusters (resolution = 0.5). These clusters show similar RNA counts, features, and percentage of mitochondrial genes captured across the three independent biological replicates indicating a high degree of consistency in our data (Figure 2a). UMAP separated by independent replicate also shows that each replicate contains cells within each cluster and that the qMcSC population in telogen skin is comparable across animals during this time point (Figure 2b). Combining the cells across samples yielded a final integrated UMAP consisting of 5545 cells arranged into eight main clusters (Figure 2c). These data are provided as a .cloupe file to view and interrogate with 10X Genomics Loupe Browser. The average *Kit* expression for each cluster ranged from 2.25-0.14 (Figure 2d-e). Substantiating this observation, we also observe variability in KIT expression at the protein level during flow cytometric analysis^1,8^.

**Figure 2:**
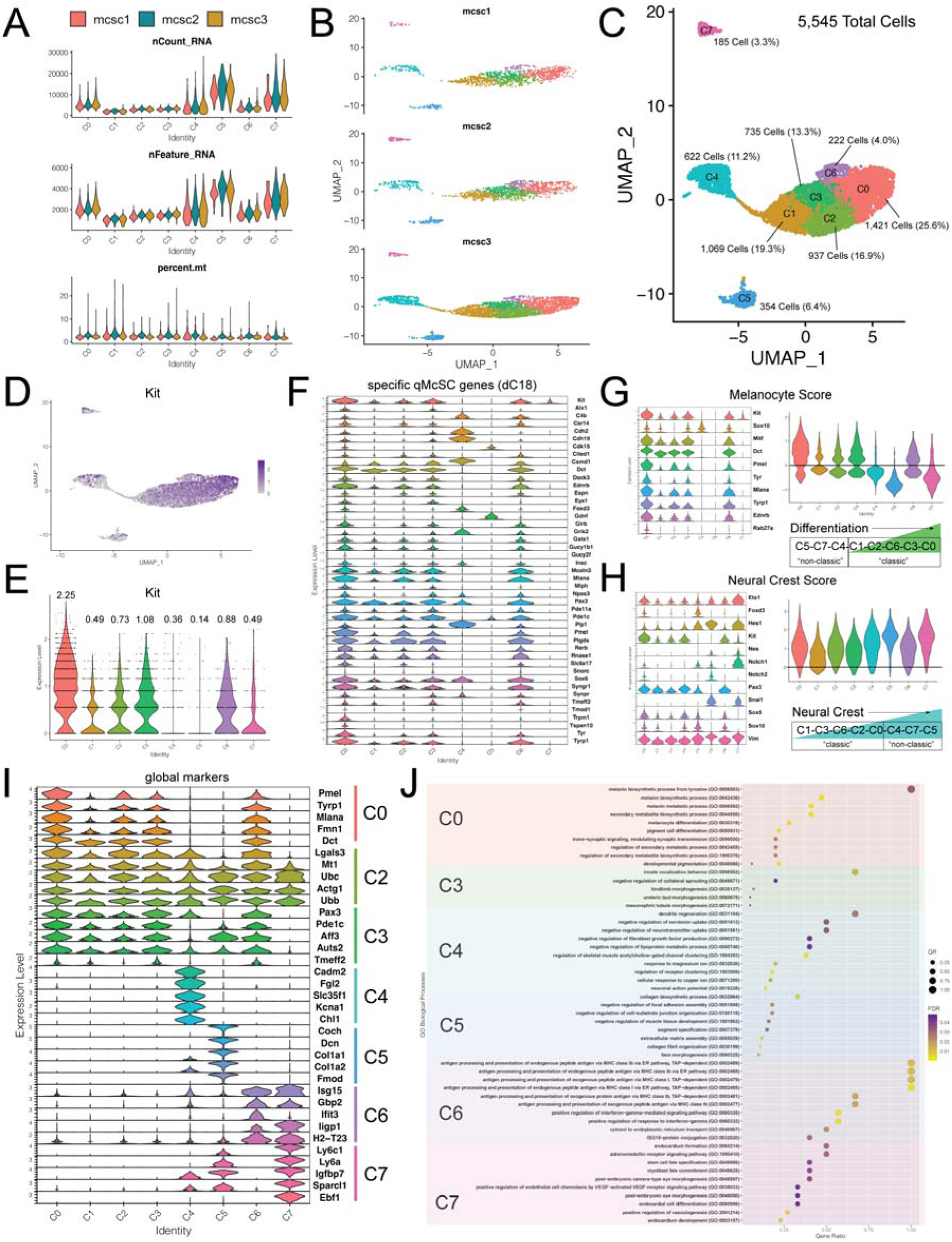
High-resolution transcriptomic map of 5,545 qMcSCs using scRNA-seq. (A) Violin plot of quality control metrics showing the average RNA, feature counts and percent mitochondrial genes detected across clusters for individual enriched qMcSC samples. (B) UMAP for each qMcSC sample showing the consistency of the clusters across replicates. (C) Combined UMAP of 5,545 *Kit+/Cd45-* cells arranged into eight subclusters, C0-C7, and the percentage of the total population for each cluster. (D-E) Plots showing variability in *Kit* expression across clusters as a UMAP (D) and violin plots (E). The numbers above each violin in (E) represent the average *Kit* expression value for the cells within the indicated cluster. (F) Violin plots showing the expression of the global markers identified for the qMcSC dC18 from the analysis of the whole dermis (see also Figure 1e) across the qMcSC clusters. (G and H) Violin plots showing the expression of melanocyte and neural crest score genes across cluster, and an accompanying violin plot showing the overall score for each cluster. The ‘Differentiation’ and ‘Neural Crest’ diagrams depict the ordering of clusters by increasing score and indicate clusters deemed “classic” and “non-classic”. (I) Violin plots showing the top five global markers with the highest average log2 fold change identified across the eight qMcSC clusters. (J) Enrichment analysis of biological processes using the top 100 global markers identified across the eight qMcSC clusters.

Using the list of specific qMcSC genes we identified in our whole dermis analysis (Figure 1e), we observe additional variability across our enriched clusters (Figure 2f). Focusing on the more distant C4, C5, and C7 clusters first, we found that these clusters had few cells represented by specific qMcSC genes and relatively low levels of melanocyte differentiation genes like *Dct*, *Tyr,* and *Pmel.* We designated C4, C5, and C7 as “non-classic” qMcSC clusters, and the remainder as “classic” qMcSC clusters. C4 shows the highest levels of cells expressing *Cdh19, Csmd1, Foxd3,* and *Plp1.* C5 is represented by cells expressing *Cdk15, Gdnf, and Pde1c.* Interestingly, the only specific qMcSC gene identified in C7 is *Kit*. C7 is also an extremely rare qMcSC subpopulation and only makes up 3.3% of the enriched KIT+/CD45-qMcSC population (Figure 2c). The ability to localize varying sets of genes to distinct clusters demonstrates the increased resolution of this data that was achieved by our sorting and filtering strategy.

To further describe all the clusters shown in Figure 2c, we scored each cluster based on melanocyte and neural crest genes (Figure 2g-h). To generate these scores, we used the AddModuleScore function in Seurat and genes associated with melanocyte differentiation and melanogenesis for the melanocyte score, and early and late neural crest transition markers for the neural crest score. We find that the highest melanocyte scores were associated with “classic” clusters 0, 1, 2, 3, and 6, and highlight a potential differentiation track with an increasing score moving from C1<C2<C6<C3<C0. Conversely, the “non-classic” clusters gave the highest values for neural crest score, increasing from C4<C7<C5, and included genes like *Hes1, Nes, Notch1,* and *Snai1*. Interestingly, C0, which has the highest “classic” melanocyte score also had the highest neural crest score of any of the “classic” clusters. Taken together we can conclude that the clusters represented in our data have characteristics consistent with the melanocytic lineage based on higher average expression of *Kit* than other cells of the dermis, expression of global markers identified in the dermis, and a high melanocyte or neural crest score.

To characterize the substructure of the enriched qMcSC pool we performed global marker detection (FindAllMarkers) across all clusters (Figure 2i, Supplemental File 3). Global marker detection will identify the top markers of each cluster based on the expression level in an individual cluster compared to the average expression across all clusters. Given that all the cells being analyzed are quiescent and have relatively low transcriptional activity, we used a logfc.threshold = 0.1 to detect global markers. Looking at the top five markers from each cluster we find C0 with highest expression of several melanogenesis genes including *Pmel*, *Tyrp1*, *Mlana*, and *Dct*, which matches this cluster’s melanocyte score, along with *Fmn1*, a gene shown to be critical for melanosome dispersion ^26^. No global markers were detected for C1. Global markers for C2 included the melanosome trafficking gene *Lgals3, Mt1*, which plays a role as an antioxidant and the detoxification of heavy metals from cells, the cytoskeleton gene Actg1, and two ubiquitination genes *Ubb* and *Ubc*. C3 is highlighted by early neural crest lineage marker *Pax3*, the phosphodiesterase *Pde1c*, the transmembrane proteoglycan *Tmeff2*, and two transcriptional activators *Auts2* and *Aff3*. C4 uniquely expressed high levels of the immune regulator *Fgl2*, a solute carrier *Slc35f1*, the potassium voltage-gated channel *Kcna1*, along with *Chl1* (L1CAM2) a neural cell adhesion molecule involved in synaptic plasticity and suppressing neuronal cell death. An additional global marker for C4 is *Cadm2*, a cell adhesion molecule we identified previously by bulk RNA-seq as upregulated in qMcSCs compared to the proliferating melanoblast precursor cells ^8^. C5 showed increased expression of *Coch* whose loss is known to cause vestibular dysfunction and hearing loss ^27^, *Dcn* which helps maintain the hair follicle niche in humans ^28^, two collagen genes *Col1a1* and *Col1a2*, and the proteoglycan *Fmod* which functions as a modulator of TGF-beta signaling ^29^. Interestingly, C6 was marked by an increase in genes related to interferon signaling including *Isg15, Gbp2*, *Ligp1*, and *Ifit3* along with the MHC class 1b gene *H2-T23*. Lastly, C7 was associated with the insulin-like growth factor regulator *Igfbp7*, *Ly6c1,* cancer and tissue-resident stem cell marker *Ly6a* (SCA-1, Stem cell antigen 1), the gene encoding the anti-proliferative and anti-tumorigenic cell matrix protein *Sparcl1* ^30^, and the transcription factor *Ebf1* that is associated with cranial neural crest development in chick embryos ^31^. The variety of global markers, along with the differences in melanocyte and neural crest score, observed across the enriched qMcSC clusters suggests different functional outcomes for each subcluster and is an idea validated by existing studies that have tested qMcSC potential (see further discussion on this topic below).

Perhaps not surprisingly, enrichment analysis of the top 100 DEGs from each cluster revealed additional heterogeneity in the biological processes associated with these clusters (Figure 2I, Supplemental File 4). No global DEGs were detected in cluster 1, and clusters C2, C3, and C6 only produced 10, 41, and 92 total DEGs, respectively. C0 was enriched for the biological processes typically associated with melanocytic cells including pigment cell differentiation (FDR = 4.57E-08, GR = 8/36, *Kit*, *Mitf*, *Ednrb*, *Tyrp1*, *Mlph*, *Slc24a5*, *Cd63*, and *Myo5a*) and melanosome organization (FDR = 1.60E-02, GR = 4/37, *Ap1s2*, *Tyrp1*, *Lyst*, and *Pmel*). No significant biological processes were enriched for the 10 DEGs identified in cluster 2. C3 DEGs were characterized as negative regulation of collateral sprouting (FDR = 4.86E-02, GR = 2/10, *Fgf13*, *Fstl4*) and developmental pigmentation (FDR = 2.96E-02, GR = 2/49, *Pax3* and *Bcl2*). The most significant biological process enriched in C4 was dendrite regeneration (FDR = 2.71E-02, GR = 2/3, *Ptn* and *Matn2*) followed by several processes associated with neuron regulation, such as neuronal action potential (FDR = 7.22E-05, GR = 6/44, *Scn7a*, *Pmp22*, *Kcna1*, *Kcna2*, *Ank3*, and *Grik2*) and regulation of long-term neuronal synaptic plasticity (FDR = 6.63E-03, GR = 4/36, *Fcgr2b*, *S100b*, *Snca*, and *Grik2*). The top processes enriched in C5 included the collagen biosynthetic process (FDR = 6.08E-03, GR = 3/9, *Col1a1*, *Serpinh1*, and *Rcn3*), negative regulation of focal adhesion assembly (FDR = 2.26E-02, GR = 3/17, *Dlc1*, *Lrp1*, *Phldb2*), and extracellular matrix assembly (FDR = 7.68E-03, GR = 4/34, *Col1a2*, *Ltbp4*, *Fbln5*, and *Gpm6b*). Positive regulation of interferon-gamma-mediated signaling pathways (FDR = 1.32E-05, GR = 4/7, *Nlrc5*, *Parp9*, *Irgm1*, and *Igtp*) was among the top processes enriched in C6 along with TAP-dependent antigen processing and presentation of endogenous peptide antigen via MHC Class 1b via ER pathway (9.56E-03, GR = 2/2, *Tap2* and *H2-T23*). Lastly, C7 was enriched for processes that included stem cell fate specification (3.81E-02, GR = 2/5, *Sox17* and *Sox18*), post-embryonic camera-type eye morphogenesis (FDR = 3.77E-02, GR = 2/5, *Kdr* and *Flt1*), and negative regulation of viral life cycle (FDR = 4.41E-03, GR = 4/34, *Ifitm2*, *Ifitm3*, *Ly6e*, and *Bst2*) (Figure 2j). Taken together these results show that the McSC population during quiescence is much more heterogeneous in gene expression and biological processes than previously acknowledged.

### Disjointed Observations of qMcSC Sub-Populations Resolved into a Single Working Model of the Population

Based on the expression of several cell-surface proteins by qMcSC subpopulations, like KIT, THY1, PLP1, CD34, and PD-L1, and the fact that some of these subpopulations have unique functional qualities, previous researchers have demonstrated that not all qMcSCs are created equal. Yet the full complexity of qMcSCs and the relationship between qMcSC subpopulations has been difficult to unify. Using the high-resolution transcriptomic landscape described above, we asked whether previously acknowledged qMcSC subpopulations could be identified and their relationship rectified within our data. We started with *Thy1* a marker for a unique population of human, interfollicular, epidermal melanocyte precursors and non-conventional, mouse melanoblasts ^32,33^. We also considered *Cd34* and *Cd274,* established markers of qMcSC subpopulations ^2,8^. C5 and C7 show exclusive and overlapping expression of *Thy1* and *Cd34*, suggesting that these markers identify the same subpopulations (Figure 3a). C6 and C7 were distinguished by high *Cd274* expression, validating our previous findings that qMcSCs physiologically express PD-L1 during telogen ^8^, and extending these findings by delineating two *Kit*+/*CD45*-/*Cd274*+ populations totaling 7.3% of the qMcSC pool (Figure 3a).

**Figure 3:**
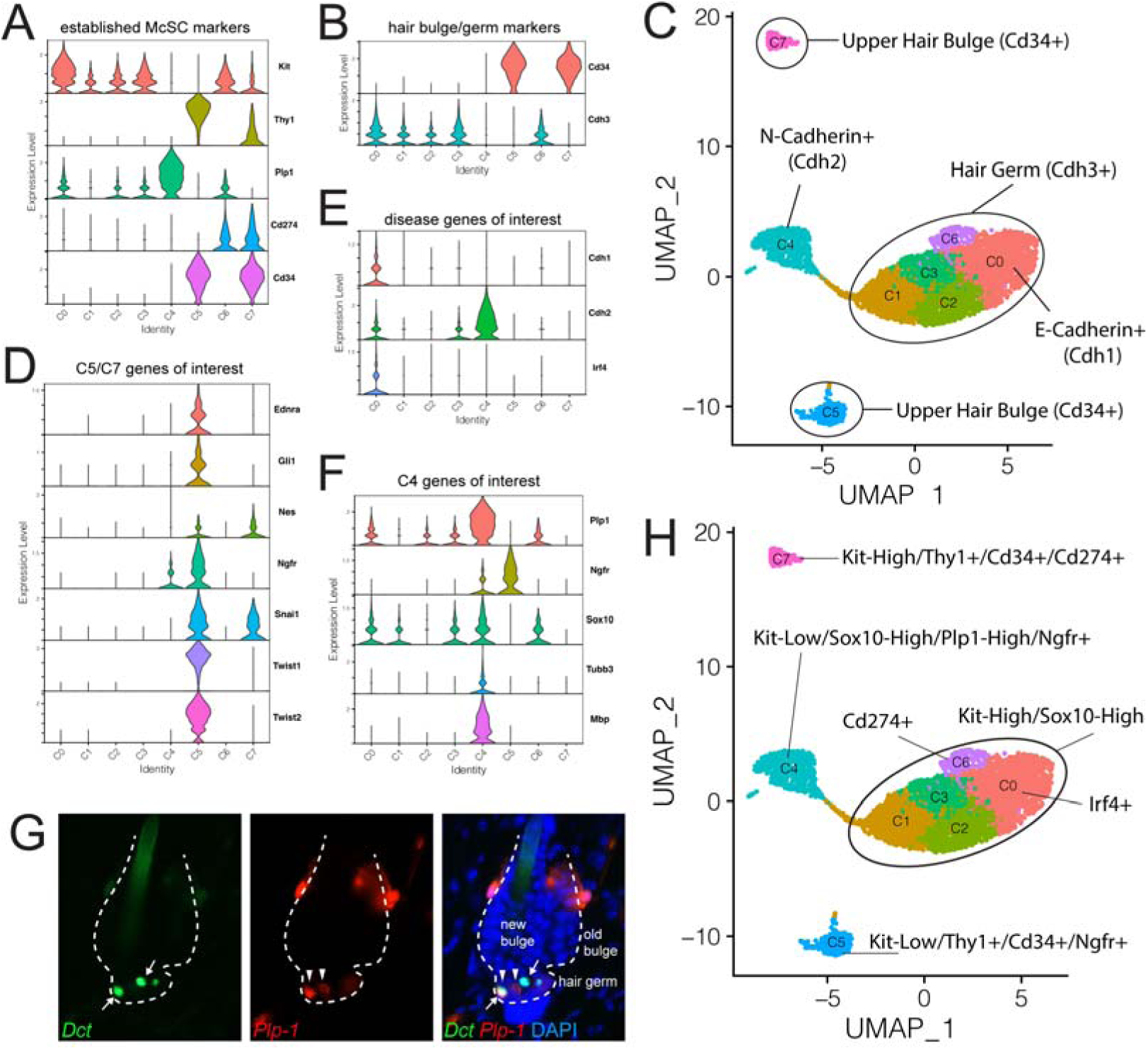
Single working model of qMcSC heterogeneity. (A) Violin plots of selected markers used previously to describe subpopulations of McSCs. (B) Violin plots of markers used to distinguish the hair bulge and hair germ. (C) UMAP depicting the predicted localization of qMcSCs within the hair follicle based on hair bulge and hair germ marker expression. (D-E) Violin plots of genes grouped as indicated. (G) Representative images of *Dct* (green), *Plp1* (red) and merged (including DAPI, blue) expression in a telogen hair of mice carrying the gene reporters *Dct-rtTA;TRE-H2B-GFP* and *Plp-CreER; Rosa-lsl-tdTomato* (general pattern observed in n=5 mice). Reporter expression was induced with doxycycline and tamoxifen, respectively, ∼2.5 weeks prior to tissue harvest. Arrows indicate *Dct*+ cells and arrowheads indicate *Plp1*+ cells localized to the the hair germ. Dotted line indicates the boundary of the hair follicle. (H) UMAP of a single working model of qMcSC subpopulation heterogeneity.

By immunohistochemistry, CDH3 (P-cadherin) and CD34 can be used to distinguish between qMcSCs that reside within two anatomically distinct regions of the hair follicle. During telogen, CD34 is restricted to the hair follicle cells and less differentiated qMcSCs localized to the upper hair bulge whereas CDH3 is associated with hair follicle cells and qMcSCs found in the lower hair germ that exhibit higher expression of the differentiation markers *Dct, Tyr, Tyrp1, Pmel,* and *Mitf* ^2,7^. By single cell RNA-seq we observe that *Cdh3* is expressed by the differentiated qMcSC clusters C0, C1, C2, C3, and C6 while *Cd34* was almost exclusively expressed by cells in C5 and C7 (Figure 3b). Accordingly, we can assign clusters anatomically with C0-C3, and C6 as hair germ McSCs and C5 and C7 as hair bulge McSCs (Figure 3c). A recent study tracking individual McSCs and their progeny using in vivo time lapse imaging showed that it is the hair germ McSCs, and not hair bulge McSCs, with the ability to give rise to differentiated bulb melanocytes ^34^. This functional difference is reiterated in our transcriptomic clustering with *Cdh3+* expressing hair germ clusters distinguishing the “classic” McSCs with higher melanocyte scores and *Cd34+* expressing hair bulge clusters marking the “non-classic” McSCs (Figure 3c and 2g). Previously, Joshi et al., 2019 discovered that CD34 also discriminates qMcSC populations with varying levels of multipotency. CD34+ qMcSCs retain the ability to differentiate into a variety of neural crest cell types whereas CD34-qMcSCs are restricted to the melanocyte lineage ^2^. In agreement with those observations, we found that the neural stem cell marker *Nes* (nestin) which is required for the proper self-renewal of neural stem cells to be exclusively expressed by C5 and C7 ^35^ (Figure 3d). We also found that the *Cd34*+ cluster C5 expresses several neural crest stem cell markers at relatively high levels including *Ednra*, *Gli1*, *Nes, Ngfr* (*p75*), *Snai1*, *Twist1*, and *Twist2*. The *Cd34*+ cluster C7 only retains expression of *Nes* and *Snai1*. This result indicates that beyond segregating qMcSCS by *Cd34* and *Cdh3* expression, *Cd34*+ qMcSCs can be further subdivided into two populations based on neural crest markers (Figure 3d), along with their differential *Kit* expression (Figure 3a).

Switching between *Cdh1* (e-cadherin) and *Cdh2* (n-cadherin) is a hallmark of the epithelial-to-mesenchymal transition and is important in both neural crest emigration from the neural tube and cancer metastasis ^36^. Perhaps surprisingly, neither *Cdh1* nor *Cdh2 is* highly represented within the qMcSC pool; *Cdh1* is only observed in C0, and *Cdh2* is most notably expressed by C4 (Figure 3e and 3c). qMcSCs are a source of cells for melanomagenesis in response to UVB exposure ^3^, and it will be interesting to test which of the qMcSC subpopulations identified here are more or less melanoma-prone or whether all qMcSCs have the same potential for cancer initiation.

Human hair graying, a phenotype attributed to McSC loss, has been genetically linked to only one gene to date, *Irf4* ^37^. Specifically, hair graying is linked to SNP rs12203592 in intron 4 of *Irf4,* the same SNP that disrupts an intronic enhancer element for TFAP2/MITF-mediated *Irf* gene transcription. IRF4 regulates *Tyr* expression cooperatively with MITF and loss of *Irf4* leads to lighter hair color in *Mitf* mutant mice ^38^. Interestingly, within our qMcSCs, *Irf4* is only apparent in C0, the most differentiated of our subpopulations (Figure 3e).

*Plp1* expression during embryogenesis has been used to lineage trace non-conventional, mouse melanoblasts derived from Schwann cell precursors. These precursors use the ventral neural crest migratory pathway and give rise to a significant number of skin melanocytes and those in extracutaneous locations like the heart and inner ear ^25,39^. *Plp1* is highest in C4 (Figure 3f), a less differentiated cell cluster by melanocyte score (compare to Figure 2g). The expression of *Plp1* by qMcSCs indicates that *Plp1* is not simply an early marker of Schwann cell precursor-derived melanoblasts but is retained in cells of the melanocyte lineage into adulthood. Validating this idea, we observe *Plp1*+ expression within the hair germ of telogen hairs from 8-week-old mice carrying both the *Dct-rtTA;TRE-GFP* ^40^ and *Plp-CreER; Rosa-lsl-tdTomato* ^41,42^ reporters. Reporter gene expression was induced postnatally in these mice with doxycycline and tamoxifen ∼2.5 weeks prior to tissue harvest. Interestingly, in the telogen hair germ we observe *Plp1+* cells that co-label with *Dct* and those that do not. This expression pattern matches that of our RNA-seq clusters with the less differentiated C4 cluster exhibiting high *Plp1* but little to no *Dct* expression, and the more differentiated C0, C2, C3 and C6 clusters exhibiting lower *Plp1* but notably higher *Dct* expression. This suggests that these *Plp1-*expressing clusters lie within a related developmental trajectory and may mark qMcSCs with varying potential. Within the C4 cluster we also observed *Ngfr+*, *Sox10+*, *Tubb3*, and *Mbp* expression (Figure 3f). Skin cells expressing *Ngfr and Sox10* are reported to retain the potential to differentiate into both melanocytes and glial cells ^43^. *Tubb3* (TUJ1) is a classic neuronal marker gene and *Mbp* expressed by both oligodendrocytes and Schwann cells. No expression of the more differentiated peripheral neuron markers neurofilament H (*Nefh*), or the peripherin gene (*Prph*) was observed ^44–46^.

In summary, these findings suggest that the heterogeneity of qMcSCs can be defined by their location within the stem cell niche, their stem and lineage potential, along with various gene expression patterns that can be unified into a single model (Figure 3c and 3h). Together these categorizations clarify the relationship between subclusters with different anatomical, molecular, and functional characteristics.

### Variable Depths of Quiescence Predicted in qMcSCs Based on Modeling the Ratio of Cell Cycle Activators to Inhibitors

Based on MCM2 protein expression and BrdU labeling, roughly 50% of the qMcSC population reenters the cell cycle during each hair cycle event ^5^. This observation suggests that qMcSCs may be held in varying states of G0 depth, with a portion of the population primed for reactivation and proliferation while others are less inclined. Similar observations have been found in other stem cell populations including muscle (G-alert vs G0), intestinal (Lgr5+/-), and hematopoietic stem cell populations ^47–49^. To determine whether a molecular signature for variable reactivation potential or quiescence depth exists in qMcSCS, we evaluated the expression of early cell cycle activators and their corresponding cell cycle inhibitors across our clusters.

Cyclins and cyclin-dependent kinases (CDKs) together are responsible for driving a quiescent cell into and through the cell cycle. To limit and control the proliferation rate, quiescent cells are frequently associated with increased expression of cyclin-dependent kinases inhibitors (CDKNs), which act as brakes to the cell cycle progression. We focused on comparing the expression of early CDKs and cyclins to the expression of corresponding CDKNs that control entry into and through the G1-phase of the cell cycle (Figure 4a-b). Initial analysis shows that *Cdk4* and *Cdk6* are highly expressed in C5 and C7. *Cdk2* has the highest expression in C0 and the lowest in C1. Assessment of the expression of the three cyclin D genes, *Ccnd1, Ccnd2, and Ccnd3*, again showed the highest expression in C5 and C7. Almost no expression of the two cyclin E genes, *Ccne1* and *Ccne2*, was observed across clusters. Next, we evaluated the expression of the *Cdkns* specifically known to interact with these early cell cycle genes. We found that inhibitors of cyclin/CDK2/CDK4 complexes including *Cdkn1a* (p21), *Cdkn1b* (p27), and *Cdkn1c* (p57) were expressed across several clusters whereas the cyclin D inhibitors including *Cdkn2a* (p16), and *Cdkn2b* (p15) were not. Interestingly, the expression of *Cdkn2d* (p19) was highly expressed by several clusters with the highest expression detected in C7. The inhibitor p19 can block the formation of the cyclin D/CDK4/CDK6 complex and arrest cells in the G0/G1 phase of the cell cycle ^50^.

**Figure 4:**
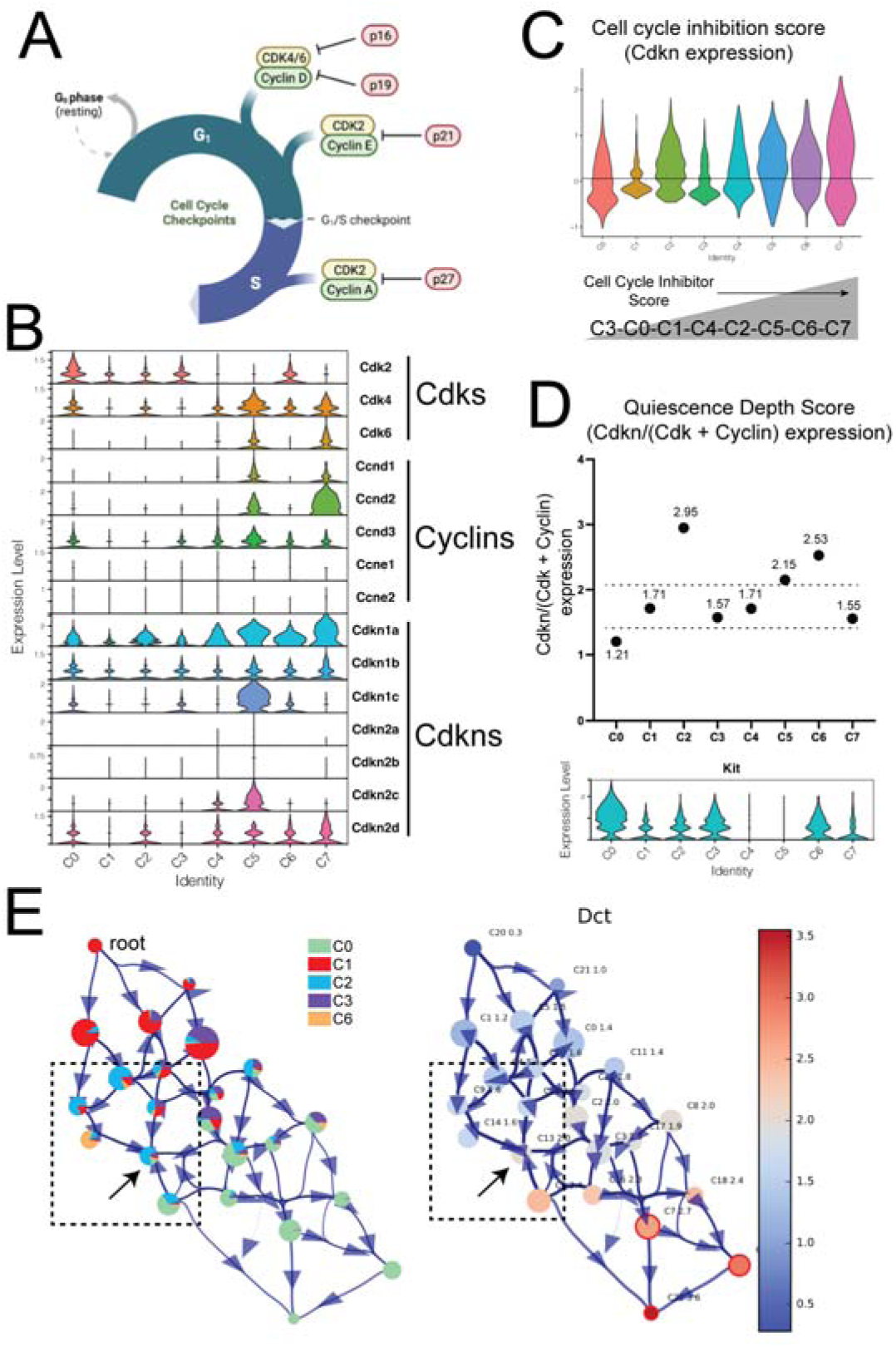
Quiescent McSCs exhibit varying levels of quiescence depth. (A) Diagram showing early cell cycle activators and inhibitors. (B) Violin plots of cell cycle activators and inhibitors across clusters. (C) Cell cycle inhibition score calculated using average expression of Cdkns. (D) The quiescence depth score calculated using the ratio of inhibitors (Cdkns) to activators (Cdks and Cyclins) showing variability in the depth of quiescence across clusters. Dotted lines indicate arbitrary boundaries between G0 states. (E) VIA trajectory model of the “classic” qMcSCs (C0-C3, C6) depicting the composition of clusters and their position along the trajectory starting from the root at C1 (left diagram) and trajectory-based expression of the melanocyte differentiation gene *Dct* (right diagram). Dotted box indicates clusters leading to a trajectory dead-end (arrow). Clusters highlighted with a red outline indicate terminal nodes.

Using the average expression of all these *Cdkns* across clusters we developed a simple “cell cycle inhibition” score to visualize the potential for cell cycle reentry across our data. We find that cell cycle inhibition scores varied across all our clusters with the highest values associated with C5, C6, and C7 and the lowest scores values associated with C0, C1, and C3 (Figure 4c). We take this one step further and use the ratio of *Cdkns* to *Cdks* and cyclin genes as a measurement of “quiescence depth” (G0 score), with higher scores indicating a decreased likelihood of reactivation (G0-deep) and lower scores indicating cells primed to reenter the cell cycle (G0-alert). Previously we showed that this ratio increases with the length of quiescence and others have shown that increased lengths of quiescence are associated with reduced rates of reactivation ^8,51^. The lowest G0 score was associated with C0 (1.21) followed by C3 and C1 (1.57, 1.71). The highest G0 scores were associated with C2, C5, and C6 (2.95,2.15, and 2.52) (Figure 4d). Of the G0-deep clusters, we characterized both C2 and C6 as “classic” qMcSCs (Figure 2g) located in the hair germ (Figure 3c), the latter of which houses the qMcSCs that transition into transit-amplifying cells to produce the differentiated hair bulb melanocytes. Interested in seeing how G0-deep status might affect the progression of these clusters towards differentiation we modeled the relationship of the “classic” qMcSCs using trajectory inference (VIA ^52^). Using C1 as the root we observe an expected trajectory that generally proceeds from C1 and ends with terminal nodes at C0. However, a number of clusters with majority C2/C6 composition appear to dead-end at the center of the trajectory (Figure 4e). Fitting this model against *Dct* gene expression, this dead-end derives from nodes that are both less and more differentiated than itself. This may reflect the McSC’s natural ability to dedifferentiate as part of the normal pigment regeneration process ^34^. In contrast, clusters with C3 composition have several edges pointing from C1 to C0. Based on these observations we predict C1, C3 and C0 as G0-alert clusters that progress developmentally in this order during the initiation of hair regrowth. We predict C2, and C6, on the other hand, are G0-deep and less likely to activate, even if located in the hair germ. The observed transcriptional heterogeneity with the qMcSC pool suggests that the choice to activate in response to a stimulus may be predefined during dormancy rather than stochastically defined at the point of stem cell activation.

Notably, C0, C1, and C3 together represent ∼58% of the qMcSC pool (see Figure 2c), which is similar to the estimated 50% of cells that undergo cell cycling during anagen ^5^. The cycling potential of qMcSCs has also been functionally attributed to dependency on KIT signaling; treating skin with the KIT function-blocking ACK2 during early anagen depletes about half the McSC pool ^5^. However, based on our transcriptional analysis, there is no clear relationship between the clusters with low G0 scores and high *Kit* expression except for C0 and C3 (compare Figure 4d and 3a). Thus, at a minimum, we anticipate C0 and C3 are the most likely qMcSC subpopulations targeted by ACK2 treatment.

### MHC Class I and Innate Immune Genes are Associated with the Two Kit+/Cd274+ qMcSC Populations

As shown in Figure 3a, two sub-populations of qMcSCs can be described as *Kit+*/*Cd274+* (C6 and C7). At the most basic level, the presence of *Cd274+* clusters in this dataset validates our previous finding that a small subpopulation of qMcSCs can be detected by surface expression of the protein product of *Cd274,* PD-L1 ^8^. This scRNA-seq expands these observations to show that there are in fact two *Kit+/Cd274+* qMcSC populations. These two cell populations can be further divided based on the expression of other markers including melanocyte differentiation markers (melanocyte score; Figure 2g), early neural crest markers (*Nes*, *Notch1, Snai1*; Figure 2h), location with stem cell niche (*Cd34+/-* and *Cdh3+/-*; Figure 3b-c), and top global markers (Figure 2i). These results show these are two distinct cell populations based on gene expression.

Despite the differences in these two *Kit+*/*Cd274+* populations, we were curious to explore similarities between these clusters that might explain their mutual upregulation of *Cd274.* Using the global markers approach (as in Figure 2i), we identified 75 global markers common to both C6 and C7 (C6/C7). Using the STRING database (string-db.org) we constructed a protein-protein interaction network to visualize the relationship between these markers. Kmeans clustering (n=4 clusters) defined two major groupings (Figure 5a), one small (18 genes) and one large (41 genes), and enrichment analysis (GO-Biological Process) revealed the small group as populated with genes involved in antigen processing and presentation (FDR = 2.8E-27, GR = 14/73) and the large group represented by genes involved in the innate immune response (FDR = 3.27E-21, GR = 22/558). Previous work on hair follicle stem cells and muscle stem cells indicates that a critical aspect of quiescence is the downregulation of MHC class molecules to evade immune detection and clearance ^53^. However, these results suggest that not all qMcSCs downregulate antigen presentation mechanisms. Our scRNA-seq data is validated by bulk RNAseq data, from both us and others ^8,10^, demonstrating that qMcSCs as a pool have significantly higher expression of MHC-I related genes (*H2-K1, H2-D1, B2m, Nlrc5*) than proliferative Mbs, activated McSCs, and McSC progeny within the hair bulb. Based on our scRNA-seq we posit that this expression comes primarily from C6/C7 and that these qMcSCs could be susceptible to immune clearance unless alternate mechanisms for immune privilege are employed. We initially identified C6 and C7 by their specific expression of *Cd274*, the gene for the immune checkpoint protein PD-L1. PD-L1, through engagement with its receptor PD-1, promotes peripheral tolerance and thus may be the mechanism to counterbalance the high potential for MHC-I presentation observed in these two *Cd274*+ clusters.

**Figure 5:**
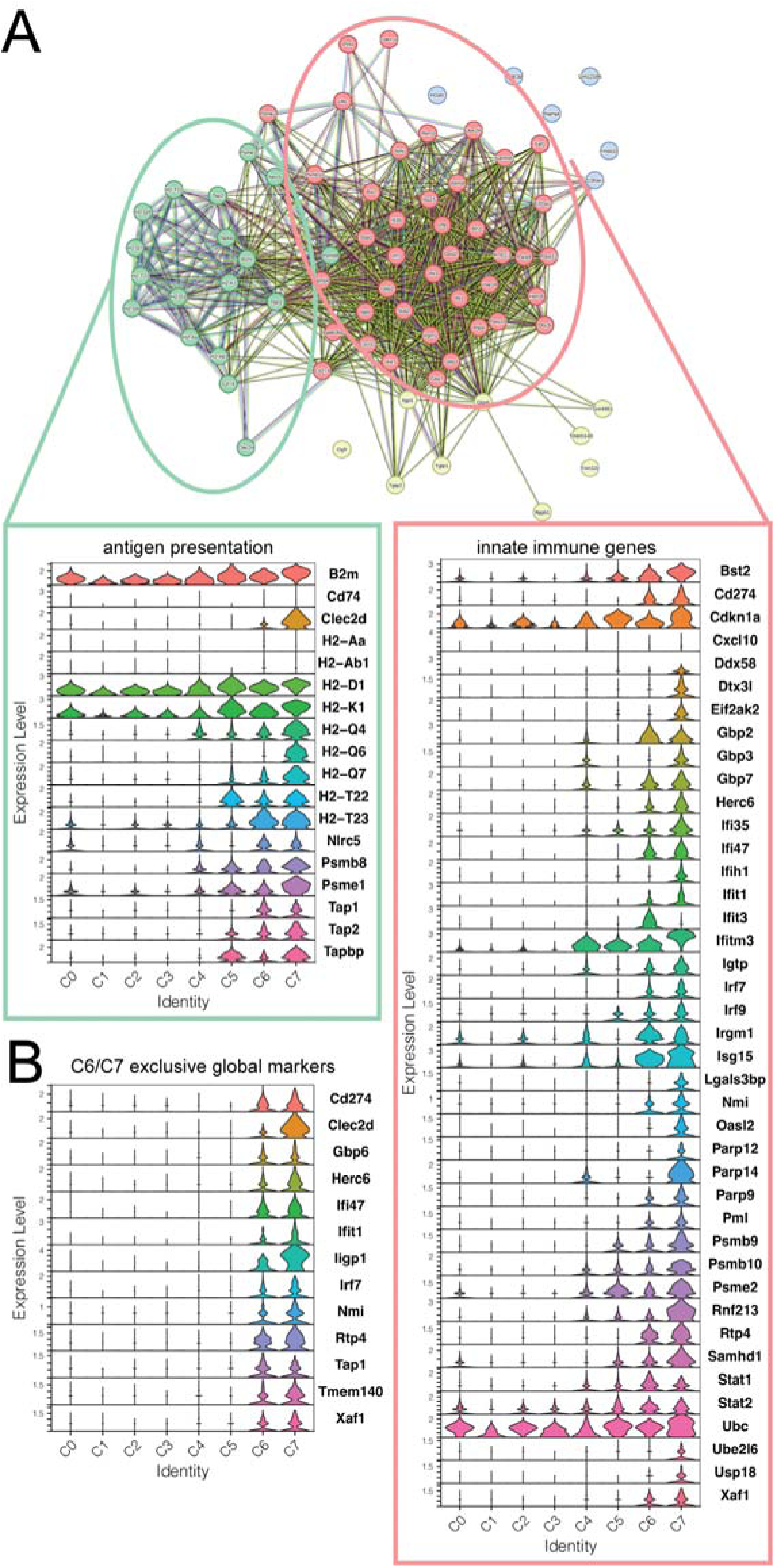
Evaluation of global markers from *Pd-l1+* clusters. (A) Protein-protein interaction (PPI) network of the overlapping global markers identified in C6 and C7 with violin plots showing the genes within each major PPI grouping. (B) Violin plot showing the 13 global markers that are specific to the *Pd-l1+* clusters C6 and C7.

Outside of *Cd274,* there are 13 other global markers that are exclusive to C6/C7 (Figure 5b). This includes the receptor transporter gene *Rtp4*, the antigen processing transporter *Tap1*, the transmembrane protein *Tmem140*, and apoptosis regulator *Xaf1*. Several genes related to the innate immune or interferon pathways mentioned above are included; *Ifi47*, *Ifit1*, the interferon-inducible GTPase *Iigp1*, *Clec2d* which can induce interferon-gamma production in human natural killer cells ^54^, and the guanylate-binding gene *Gbp6* which is induced by interferon ^55^. We also observe, and the ubiquitin ligase *Herc6* ^56^, *Nmi* which is known to regulate the innate immune response by interacting with STAT and MYC proteins ^57^, and the interferon regulatory factor *Irf7*. The latter genes, *Nmi* and *Irf7,* are of particular interest because they provide a potential transcriptional mechanism for the specific expression of *Cd274* in C6/C7. IRF7 can bind directly to the promotor of *Cd274* and enhance its expression in both human and mouse cell lines independent of the canonical signal, interferon-gamma ^58^ and *Nmi,* also known as N-myc and STAT interactor, is known to regulate the innate immune response by interacting directly with STAT1 and STAT5 ^57,59^. *Stat1* and *Stat2* are also global markers common to C6/C7 and whose proteins participate in *Cd274* activation ^60,61^. Altogether, the unique expression of *Cd274* in C6/C7 may be the consequence of these upstream signaling and transcriptional cascades that are also highly expressed in C6/C7.

## Discussion

Until this study, investigating the heterogeneity of qMcSCs remained difficult due to the rarity of this stem cell population. Using enrichment methods to isolate KIT+/CD45-cells of the dermis we were able to generate a high-resolution transcriptomic map of qMcSCs that reconciles previously reported subpopulations and highlights novel qMcSC subpopulations. Across these qMcSC clusters, we detected differences in differentiation states, quiescence depth, and immune status. The results of this study provide a new working model of the qMcSC population and a baseline for future experiments focused on evaluating population dynamics within qMcSCs under perturbation.

One example of how this data could be employed to generate new hypotheses comes from our novel observations regarding C6/C7. The immune genes that define the common global markers of C6/C7 are reminiscent of a similar signature we observed in qMcSCs in *Mitf^mi-vga^*^9^*^/+^* mice using bulk RNA-seq ^1^. In fact, 33 of the 74 global markers from C6/C7 are also DEGs upregulated with *Mitf* haploinsufficiency. *Mitf* knockdown and ChIP suggest that MITF can transcriptionally repress several interferon-stimulated genes (ISG) observed in C6/C7 including *B2M, H2-T23, Ifih1, Ifit3, Isg15, Stat1,* and *Tap1.* The presence of unique subpopulations of qMcSCs that express these genes physiologically in wildtype animals could suggest that upregulation of these genes in *Mitf^mi-vga^*^9^*^/+^* mice may not simply reflect a gain-of-function transcriptional reprogramming but rather a transition of the qMcSC pool towards an existing C6/C7 state. Alternatively, the increase in the expression ISGs observed in *Mitf^mi-vga^*^9^*^/+^* mice could result from differences in the composition of the qMcSC population as a whole with a decrease in the percentage of more differentiated clusters like C0 thus making the entire qMcSC population in bulk RNA-seq appear to be expressing higher levels of ISGs compared to wild-type animals. Both of these possibilities should be considered when designing future experiments and employing scRNA-seq analysis will be able to directly test both of these hypotheses. It will be interesting to consider these scRNA-seq data in the context of other bulk sequencing datasets comparing McSCs under different conditions (e.g., ionizing radiation, melanoma formation following UVB exposure, or depletion with age). In summary, this detailed evaluation of the qMcSC pool using scRNA-seq provides new avenues for investigating McSC characteristics and regenerative potential.

## Materials and Methods

### Animals

All C57BL/6J female mice used in this study for single cell RNA-seq analysis were obtained directly from JAX (at 8 weeks of age) and housed in standard cages with a 12-hour light/dark cycle for a minimum of one week to allow for adjustment before use.

### Fluorescent-activated cell sorting

Whole skin was harvested from nine-week-old mice and processed for single-cell suspension using methods previously described ^1,8^. Briefly, cell suspensions were labeled with anti-KIT and anti-CD45 antibodies and sorted into 1.5ml tubes containing 1mL of 10% FBS to reduce cell loss during collection. Cells were then spun at 200g for 5 mins, the supernatant was carefully removed, and 30ul of BSA was added to resuspend the pellet and placed on the ice during short transport to the sequencing facility.

### Alignment and Quality Control of Sequencing Data

All sequencing files were aligned to the mm10 transcriptome using Cell Ranger (v5.0.1). on the UAB high-performance cluster computer (Cheaha). Following alignment barcodes, features, and matrix files were loaded in R and a single-cell object was created using Seurat (v4.0.4, min.cells = 5, min. features = 100). The percentage of the mitochondrial and percent largest gene was then calculated and stored within the Seurat object. Quality metrics were then individually assessed and cells more than three standard deviations from the mean of the metric were then removed by sub-setting the data. Additionally, cells were only considered if they had greater than 100 features and an RNA count above 50 were removed (McSC1 = 205, McSC2 = 147, McSC3 = 345, Derm = 191). The data was then normalized across features using the centered log ratio (CLR) method before scaling using default parameters. Variable features were then detected using the VST selection method across the mean number of features detected in cells from each sample followed by dimension reduction. Following principal component analysis, nearest neighbors (dims = 1:20) and clusters (resolution = 1.0 for dermis) were identified prior to generating UMAP (dims = 1:20) and saving the object as an RDS file.

Individual McSC files were filtered as described above prior to merging. The combined object was then normalized across features using the CRT method with default scaling. Variable features of the merged object were then detected using VST selection method across the mean number of features detected across cells (nfeatures = 2588). Dimension reduction was performed similarly as described above with principal component analysis followed by nearest neighbors (1:20) and clustering (resolution = 0.3) followed by generation of UMAP (dims = 1:20). The merged object was then further filtered to removed clusters with low expression of Kit (< 0.1) and high expression of hair follicle markers Krt14 and Krt15. Variable features (n = 3000), nearest neighbors (dims = 1:20), clustering (resolution = 0.3), and UMAP generation (dims = 1:20) was performed a final time on the remaining 5545 cells. Global differential expression was determined on the dermis and McSC Seurat objects by using the function “FindAllMarkers”. Neural crest, melanocyte, and CDKN scores were calculated using the “AddModuleScore” and the gene lists specified in the text. Quiescence depth scores were calculated by combining the average expression of all Cdkns and dividing by the combined total of Cdks and Cyclins with the final plot being generated using Prism. Enrichment analysis of biological processes and cellular components was performed using the Panther database. Raw sequence data is available at NCBI GEO (GSE# XXXXX). The filtered (5545 cells) McSC Seurat object used to generate the UMAPs in Figures 2 and 3 is also available as a .cloupe file (Supplemental File 2) to view and interrogate with Loupe Browser (10X Genomics).

### VIA trajectory analysis

For the exploration of cellular dynamics and gene expression patterns within the selected clusters the Seurat object from the above-mentioned section was converted to the loom object using the loomR R package. The resulting loom file and other supporting files were imported into the Python (v3.8.18) environment for further analysis using the Scanpy (v1.9.6) package as an AnnData object. The “C1” cluster was set as root (root_user), ncomps as 30, and the knn parameter was set to 10 for the VIA analysis using the pyVIA (v0.1.96) package. The cluster composition and pseudotime diagrams were generated using the plot_piechart_viagraph function. The gene expression along the VIA graph was plotted using the plot_viagraph function.

### Assessment of Plp1 and Dct co-expression

Secondary analysis of tissues obtained previously from mice carrying *Dct-rtTA;TRE-H2B-GFP* (NCI-#01XT4) ^1^ and *Plp-CreER* (JAX, #005975)*; Rosa-lsl-tdTomato* (JAX, #007908) reporter genes was used to assess *Plp1* and *Dct* co-expression within the hair follicle. *Dct-rtTA;TRE-H2B-*GFP, *Plp-CreER*, and *Rosa-lsl-tdTomato* were obtained from the National Cancer Institute (NCI) and the Jackson Laboratories (JAX), respectively, and maintained by the Center of Animal Resources and Education (CARE) at Cornell University College of Veterinary Medicine. Animal genotypes were determined by PCR following the protocols provided by the Jackson Laboratories. Mice were housed in standard cages with a 12-hour light/dark cycle. At 8 weeks of age, during the dormant telogen hair cycle, mice were induced for reporter gene expression. From day 1 to day 5, mice were treated with doxycycline (Alfa Aesar, Cat#J60579; 200mg/L) in their drinking water and given 200ul tamoxifen daily (Cayman, Cat#13258; 10mg/ml in 10% EtOH/90% corn oil) by intraperitoneal injection. For primary experimental purposes unrelated to these secondary analyses, these mice were also irradiated with 2.2 J/m^2 UVB once every other day for three exposures. UVB activation of qMcSCs will induce their migration to the epidermis and these mice were generated to track *Plp1* and *Dct* expression in qMcSCs prior to their relocation. Dorsal skin was collected at 7 days post third UVB. Skin was fixed in formalin overnight at 4°C and embedded in OCT (Fisher, Cat#23730571) blocks. Tissue blocks were sectioned at 8µm thickness for imaging. Sections were fixed in formalin for 10 mins, followed by two H2O washes (10mins each). Sections were mounted with Fluoroshield with DAPI (Abcam, ab104139). Images were taken using the Leica DM7200 fluorescence imaging platform with LAS X version 3.7.5.

## Acknowledgements

We thank Drs. Stephanie Dickinson and Andrew Brown (UAB Nathan Shock Center Data Analytics Core and Indiana University Bloomington) for their early advice on statistical considerations of single cell datasets. We also appreciate Dr. George Green (UAB) for lending his technical skills with Seurat and Loupe Browser.

